# Hypermetabolic state is associated with circadian rhythm disruption in mouse and human cancer cells

**DOI:** 10.1101/2023.11.08.566310

**Authors:** Daniel Maxim Iascone, Xue Zhang, Patricia Bafford, Clementina Mesaros, Yogev Sela, Samuel Hofbauer, Shirley L. Zhang, Kieona Cook, Pavel Pivarshev, Ben Z. Stanger, Stewart Anderson, Chi V. Dang, Amita Sehgal

## Abstract

Crosstalk between cellular metabolism and circadian rhythms is a fundamental building block of multicellular life, and disruption of this reciprocal communication could be relevant to degenerative disease, including cancer. Here, we investigated whether maintenance of circadian rhythms depends upon specific metabolic pathways, particularly in the context of cancer. We found that in adult mouse fibroblasts, ATP levels were a major contributor to overall levels of a clock gene luciferase reporter, although not necessarily to the strength of circadian cycling. In contrast, we identified significant metabolic control of circadian function in an *in vitro* mouse model of pancreatic adenocarcinoma. Metabolic profiling of a library of congenic tumor cell clones revealed significant differences in levels of lactate, pyruvate, ATP, and other crucial metabolites that we used to identify candidate clones with which to generate circadian reporter lines. Despite the shared genetic background of the clones, we observed diverse circadian profiles among these lines that varied with their metabolic phenotype: the most hypometabolic line had the strongest circadian rhythms while the most hypermetabolic line had the weakest rhythms. Treatment of these tumor cell lines with bezafibrate, a peroxisome proliferator-activated receptor (PPAR) agonist shown to increase OxPhos, decreased the amplitude of circadian oscillation in a subset of tumor cell lines. Strikingly, treatment with the Complex I antagonist rotenone enhanced circadian rhythms only in the tumor cell line in which glycolysis was also low, thereby establishing a hypometabolic state. We further analyzed metabolic and circadian phenotypes across a panel of human patient-derived melanoma cell lines and observed a significant negative association between metabolic activity and circadian cycling strength. Together, these findings suggest that metabolic heterogeneity in cancer directly contributes to circadian function, and that high levels of glycolysis or OxPhos independently disrupt circadian rhythms in these cells.

## Introduction

The circadian clock coordinates the timing of key physiological processes in mammals by controlling the expression of rate-limiting enzymes in diverse metabolic pathways [1, 2]. The cellular clock itself is driven by transcription/translation feedback loops composed of a set of transcriptional activators and repressors [3, 4]. The major activators BMAL1 and CLOCK form a heterodimer to induce transcription at E-box enhancer elements of many genes, including their negative regulators Period (PER) and Cryptochrome (CRY); feedback by PER-CRY results in 24-hour rhythms of gene expression [5]. In turn, clock-controlled processes like glycolytic and mitochondrial metabolism regulate circadian rhythms through their use and production of metabolites such as NAD^+^, acetyl-CoA, and reactive oxygen species that act as co-factors and modifiers of clock-associated proteins [6-9].

Crosstalk between basic cellular physiology and circadian rhythms is essential for the normal function of healthy adult cells, but in disease states this mutual regulation may produce a feedback loop that drives progressive dysfunction. For example, clinical data reveal this connection between the circadian clock and cancer-not only can circadian misalignment such as chronic shiftwork contribute to oncogenesis, but cancer progression itself is capable of further disrupting the circadian clock [10-13]. However, many cancers retain clock function, raising the question of what it is in different cancerous cell types that sustains or disrupts circadian rhythms.

Given that cancers tend to undergo metabolic reprogramming, we considered the possibility that maintenance of circadian rhythms depends upon a specific metabolic pathway and is lost when cells switch to relying on a different pathway. In the present study, we investigated the contribution of glycolysis and OxPhos toward the regulation of circadian rhythms across a panel of cell lines derived from a mouse model of pancreatic adenocarcinoma (PDA). We observed considerable diversity in both the metabolic and circadian phenotypes among these lines that suggest a strong inverse association between levels of glycolysis/OxPhos and the strength of circadian cycling. Enhancing OxPhos in these cell lines weakened circadian cycling, while inhibiting OxPhos strengthened circadian cycling only in the single line that began with low levels of glycolysis. In addition, across a panel of human patient-derived melanoma cell lines, we observed a significant negative association between metabolic activity and circadian cycling strength such that lines with the highest levels of ATP production and basal glycolysis demonstrated relatively weak circadian oscillation. Together these data suggest a model in which circadian cycling in cancer is enhanced by a hypometabolic state and weakened by hypermetabolic state. Our data demonstrate direct and independent control of the molecular clock by both glycolytic and oxidative metabolism in mouse and human cancer models.

## Results

### Luciferase reporters as a model system for metabolic-circadian crosstalk

To determine the role of oxidative phosphorylation (OxPhos) in regulating the circadian clock, we established a model system of pharmacological OxPhos modulation in mouse adult fibroblasts (MAFs) expressing the circadian luminescent reporter Bmal1::luciferase [14]. We chose bezafibrate (BEZ), a PPAR agonist, to enhance OxPhos without activating glycolysis by stimulating acetyl-CoA production through fatty acid oxidation (**Fig. 1A-C**) [15, 16]. We inhibited OxPhos in these cells with the potent mitochondrial complex I inhibitor rotenone (ROT) [17]. To validate the use of these drugs to modulate OxPhos we used the Seahorse mitochondrial stress assay in which cellular oxygen consumption associated with ATP production is measured as the difference between baseline oxygen consumption rate (OCR) and OCR following injection of the ATP synthase inhibitor oligomycin [18]. BEZ and ROT treatment are sufficient to alter both MAF mitochondrial ATP production and maximum respiration capacity (the difference between OCR after treatment with mitochondrial membrane uncoupler FCCP and non-mitochondrial OCR) (**Fig. 1B-C**).

**Figure 1.**
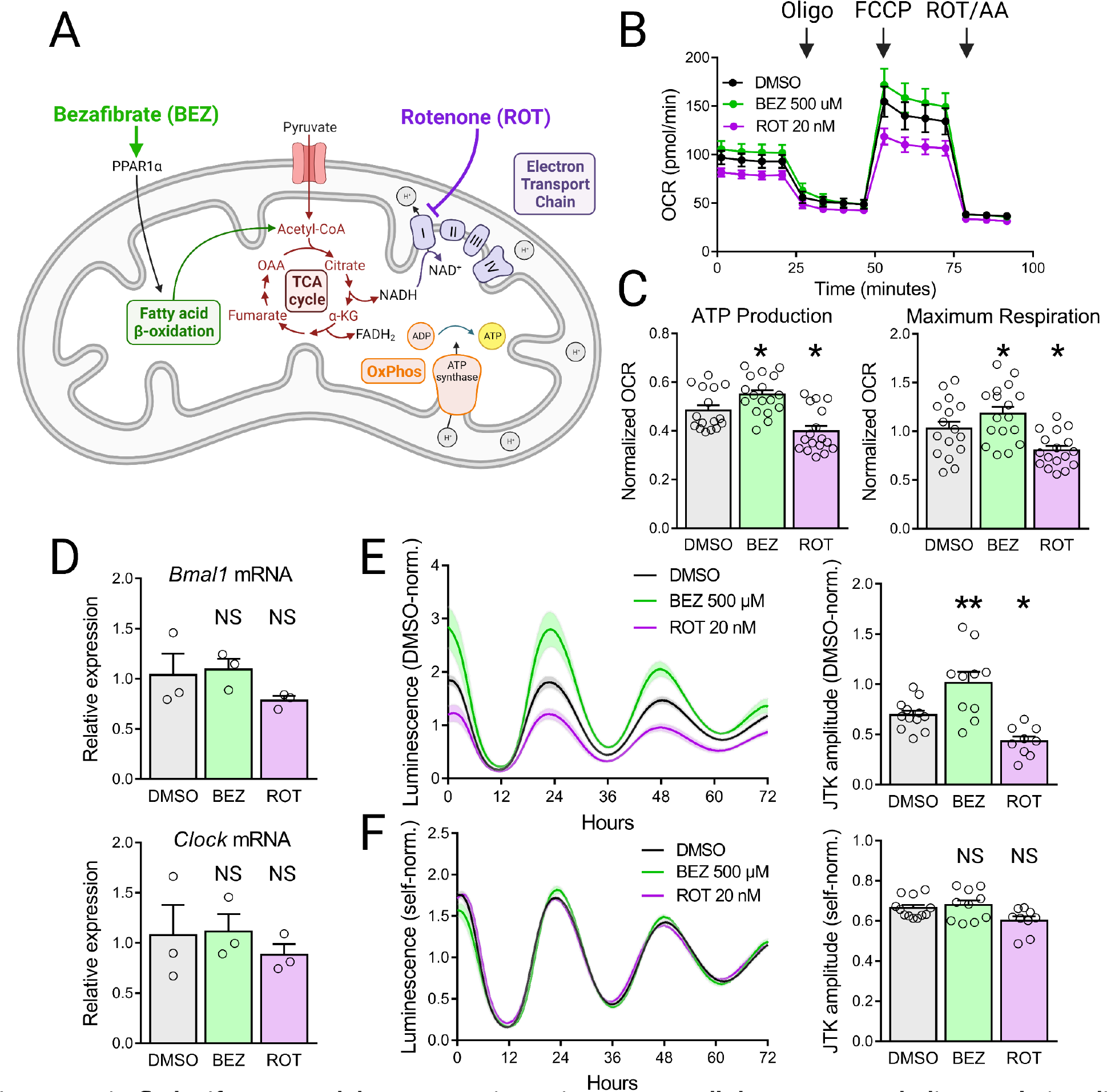
Firefly luciferase model system to investigate crosstalk between metabolism and circadian rhythms. (**A**) Schematic illustration of pharmacological modulation of OxPhos. Bezafibrate (BEZ) enhances OxPhos by driving fatty acid oxidation through PPAR agonism, increasing the production of acetyl-CoA. Rotenone (ROT) reduces OxPhos by directly inhibiting Complex I of the electron transport chain. (**B**) Oxygen consumption rate (OCR) of mouse adult fibroblasts (MAFs) treated with BEZ and ROT for 48 hours prior to a Seahorse mitochondrial stress test (n = 3 experiments). (**C**) ATP production (left) and maximum respiration capacity (right) of MAFs calculated from the OCR traces in (B). (**D**) qPCR measurement of peak *Bmal1* (top) and *Clock* (bottom) expression relative to *Actin* of MAFs treated with BEZ and ROT for 48 hours (n = 3 biological replicates) (**E**) DMSO condition-normalized luminescence traces (left) and oscillation amplitudes (right) recorded from Bmal1::luciferase MAFs treated with BEZ and ROT reflect combined changes in ATP production and circadian gene expression (n = 3 experiments) (**F**) Self-normalized luminescence traces (left) and oscillation amplitudes (right) reflect circadian gene expression alone (n = 3 experiments). Error bars indicate mean ± SEM. NS not significant, ∗p < 0.05, ∗∗p < 0.01, one-way ANOVA test with Benjamini, Krieger, and Yekutieli’s two-stage step-up procedure to control the FDR, drug treatment conditions compared to DMSO control.

Despite significant changes in the drug-treated MAF circadian luminescence signal relative to the control condition, however, surprisingly we detected no changes in peak *Bmal1* and *Clock* mRNA expression in MAFs following 48 hours of either BEZ or ROT treatment (**Fig. 1D-E** and see **Fig. S2**). We believe that this apparent inconsistency is due to the reliance of the luciferase reporter enzyme on ATP to catalyze the luminescence-generating cleavage of luciferin [19, 20]. Therefore, any analysis of luciferase-based circadian reporters that preserves changes in the magnitude of the luminescence signal (such as normalizing drug-treated samples to a control condition) would represent effects of both circadian patterns of transcription and intracellular ATP levels. Crucially, we were able to isolate the transcriptional component of the luminescence signal by normalizing time points within each trace only to their own 24-hour mean value. This self-normalization effectively cancels out differences among conditions associated with the ATP-dependent signal magnitude and leaves only the underlying oscillation driven by the circadian promoter fragment. Consistent with their clock gene expression patterns, self-normalized Bmal1::luciferase MAF traces show no effect of BEZ or ROT treatment on JTK amplitude (**Fig. 1F**).

To confirm that self-normalization of circadian luciferase traces is able to identify changes related to clock gene transcriptional activity, we analyzed MAFs treated with Nrf2 agonist dimethyl fumarate (DMF)-a manipulation shown to increase oxygen consumption of human fibroblasts as well as dampen circadian cycling by driving expression of negative clock regulators *Cry2* and *Rev-erbα* [21, 22]. As expected, we observed a significant decrease in the oscillation amplitude of self-normalized Bmal1::luciferase traces from MAFs treated with DMF (**Fig. S1**).

To rule out the contribution of the cell cycle or cell viability to circadian rhythms in these cells, we determined there to be no significant differences in levels of cell division or apoptosis in MAFs treated with BEZ or ROT relative to control conditions (**Fig. S3-4**). Together, these results establish a paradigm for investigating crosstalk between oxidative metabolism and circadian rhythms with luciferase reporters.

### Metabolic profiling of PDA cell lines

We next chose to explore the role of metabolism in the regulation of circadian rhythms in the context of cancer, in which disease-associated metabolic reprogramming is associated with both tumor development and malignant progression [23]. This reprogramming, driven by the cellular response to inflammation and the hypoxic tumor microenvironment, includes transcriptional programs that enhance glycolysis while inhibiting OxPhos, by increasing glucose uptake, upregulating rate-limiting glycolytic enzymes, and driving the conversion of pyruvate to lactate (**Fig. 2A**) [24]. To isolate the contribution of metabolism to circadian regulation independent of genetic variability, we generated circadian reporter cell lines from a library of congenic tumor cell clones derived from a mouse model of pancreatic adenocarcinoma (PDA) induced with KRAS and p53 mutations [25]. We first performed metabolic profiling of these clones using liquid chromatography-high resolution mass spectrometry (LC-HRMS) to identify cell lines with diverse metabolic phenotypes (**Fig. 2B**). We used relative levels of pyruvate to lactate to estimate glycolytic and oxidative metabolic activity in these lines. Conversion of pyruvate to acetyl-CoA drives the production of electron donors by the TCA cycle necessary for the function of the electron transport chain and OxPhos. In contrast, cancer-associated metabolic reprogramming stimulates glycolysis and the conversion of pyruvate to lactate such that a low pyruvate/lactate ratio would indicate elevated reliance on glycolysis. The PDA cell lines we chose for this study represent four distinct metabolic phenotypes: a hypometabolic line (6419) with low levels of pyruvate and lactate, a hypermetabolic line (6499) with high levels of pyruvate and lactate, a glycolytic line (6556) with high levels of lactate relative to pyruvate, and a line with high levels of OxPhos (2699) based on high levels of pyruvate relative to lactate. Relative ATP levels among these lines also supported this categorization as the hypometabolic line 6419 demonstrated the lowest ATP levels and the hypermetabolic line 6499 demonstrated the highest ATP levels (**Fig. 2B**).

**Figure 2.**
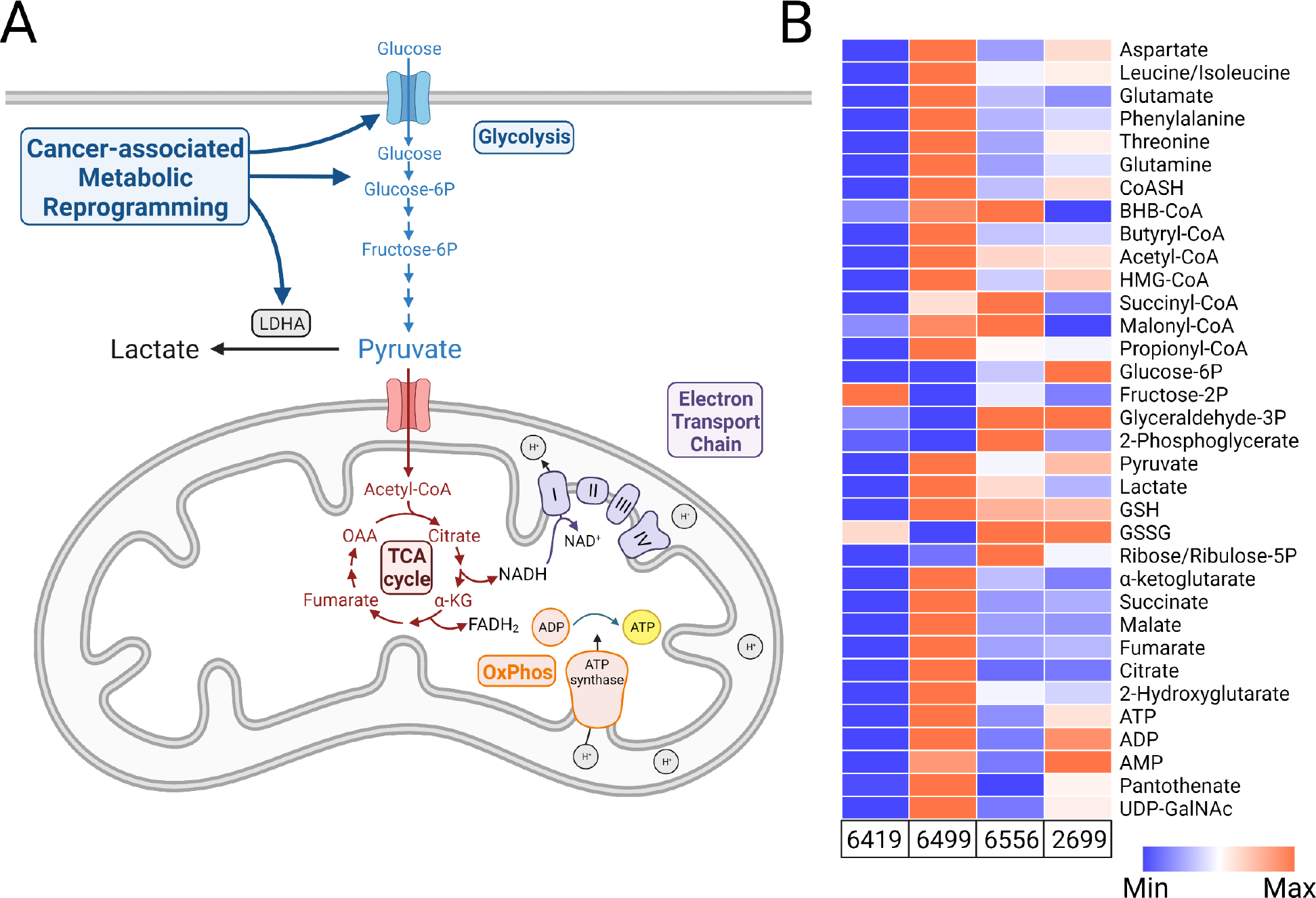
PDA cell lines have diverse metabolic phenotypes. (**A**) Schematic illustration of cancer-associated metabolic reprogramming. During cancer progression, the hypoxic environment of the tumor stimulates transcriptional programs that enhance glycolysis while inhibiting OxPhos by increasing glucose uptake, upregulating rate-limiting glycolytic enzymes, and driving the conversion of pyruvate to lactate. (**B**) Metabolite profiling with liquid chromatography coupled to high-resolution mass spectrometry (LC-HRMS) reveals significant metabolic diversity among the PDA clones chosen to generate circadian reporter lines.

### PDA circadian cycling strength varies with metabolic state

To investigate the association between metabolic activity and circadian rhythms in PDA, we generated circadian reporter lines through lentiviral transfection of Per2::luciferase. Strikingly, the hypometabolic cell line 6419 had the strongest circadian amplitude while the hypermetabolic line 6499 had the weakest rhythms, suggesting a paradigm in which overall metabolic activity inhibits circadian cycling in PDA (**Fig. 3A-B**). Consistent with this interpretation, the glycolytic line 6556 and the high OxPhos line 2699 demonstrated intermediate circadian cycling strengths (**Fig. 3C-D, Fig. 3E** top panel). Notably, the glycolytic line 6556 demonstrated significantly weaker circadian cycling than the high OxPhos line 2699, suggesting that cancer-associated metabolic reprogramming toward glycolytic metabolism has a dampening effect on circadian rhythms. Despite significant differences in circadian amplitude among all PDA cell lines relative to one another, we detected no changes in period length among these lines suggesting that metabolic diversity in this context contributes principally to circadian cycling amplitude (**Fig. 3E** bottom panel).

**Figure 3.**
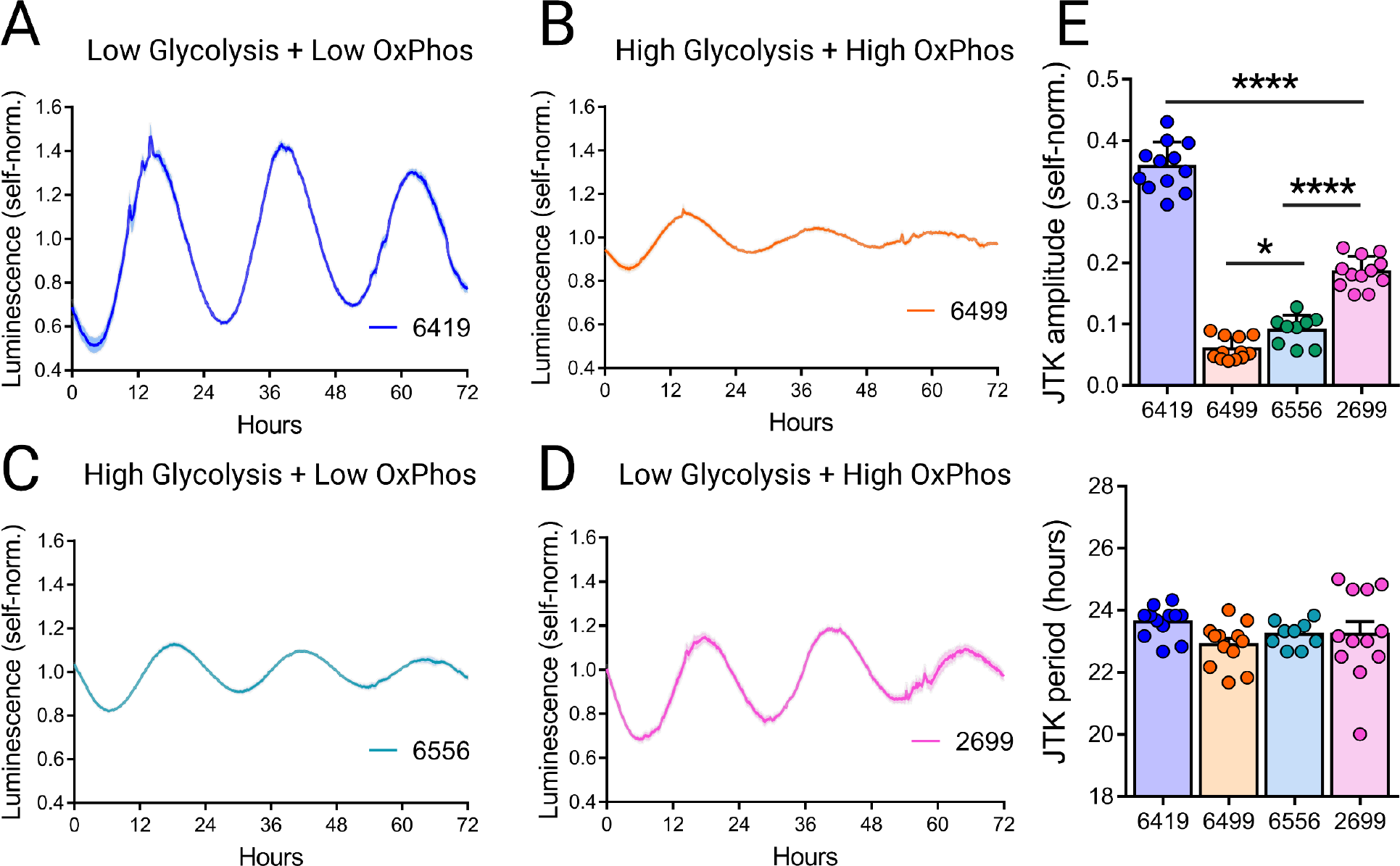
PDA cell lines have diverse circadian phenotypes. (**A-D**) Self-normalized luminescence traces ± SEM of Per2::luciferase activity recorded from PDA cell lines 6419 (A), 6499 (B), 6556 (C), and 2699 (D) (n = 3 experiments). (**F**) Oscillation amplitudes (top) and periods (bottom) calculated with JTK analysis of the traces in A-D. Error bars indicate mean ± SEM. ∗p < 0.05, ∗∗∗∗p < 0.0001, one-way ANOVA test with Holm-Sidak’s multiple comparisons test.

### Pharmacological inhibition of OxPhos enhances PDA circadian cycling only when paired with low glycolysis

To establish a causal relationship between metabolic activity and circadian regulation in PDA, we next assessed whether pharmacological modulation of OxPhos could alter circadian cycling in these lines. Notably, treatment with BEZ or ROT was sufficient to alter ATP production in these cell lines as measured by a shift in luciferase signal magnitude in the drug-treated conditions relative to controls (**Fig. 4A-D** left panel). 500 μM BEZ treatment was sufficient to increase luciferase activity-driven luminescence in all PDA lines, with the strongest effect in lines 6419 and 6556 that demonstrated the lowest levels of OxPhos prior to treatment. As expected, BEZ treatment had the weakest effect in the hypermetabolic line 6499 that previously demonstrated the highest ATP levels. 20 nM ROT treatment decreased luminescence in all PDA lines except for the hypometabolic line 6419 that demonstrated the lowest ATP levels among all cell lines. Together these results suggest that BEZ and ROT treatment are sufficient to modulate OxPhos in PDA cells similarly to MAFs except for possible ceiling/floor effects in the hypo- and hypermetabolic PDA lines.

**Figure 4.**
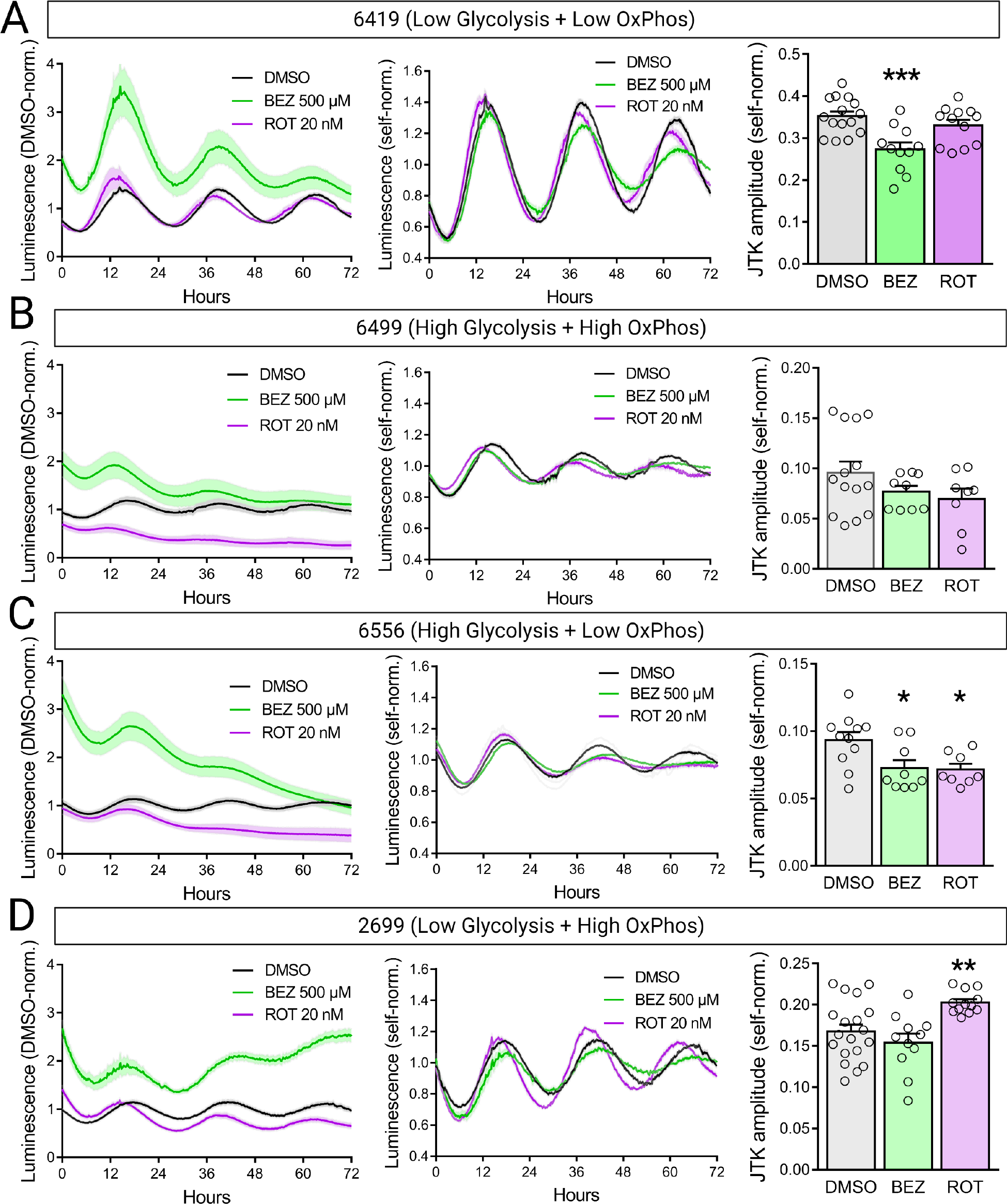
Pharmacological modulation of OxPhos in PDA cell lines. (**A-D**) DMSO condition-normalized Per2::luciferase luminescence traces (left), self-normalized traces (middle), and self-normalized oscillation amplitudes (right) recorded from PDA cell lines 6419 (A), 6499 (B), 6556 (C), and 2699 (D) (n = 3 experiments). Error bars indicate mean ± SEM. ∗p < 0.05, ∗∗p < 0.01, ∗∗∗p < 0.001, one-way ANOVA test with Benjamini, Krieger, and Yekutieli’s two-stage step-up procedure to control the FDR, drug treatment conditions compared to DMSO control.

Considering that the hypermetabolic PDA line 6499 demonstrated the weakest circadian rhythms prior to treatment, we hypothesized that enhancing OxPhos in this line would inhibit circadian cycling. Interestingly, BEZ treatment significantly weakened circadian cycling only in PDA lines 6419 and 6556, the two lines that demonstrated the greatest increase in ATP production after BEZ treatment (**Fig. 4A** and **4C**). In contrast, inhibition of OxPhos with ROT treatment was sufficient to enhance circadian cycling only in a single PDA line, 2699, whose metabolic profile suggested low glycolysis and high OxPhos prior to treatment (**Fig. 4D**). As inhibition of OxPhos in this line would establish low levels of both glycolysis and OxPhos, enhancement of circadian cycling in this condition further strengthens the association between hypometabolic state and circadian rhythmicity we observe in this PDA model. Despite some differences in cell death after 48 hours of BEZ or ROT treatment across PDA lines, overall apoptosis levels in all conditions remained quite low (<5% of total cells) in PDA cultures suggesting that apoptosis had minimal or no effect on observed differences in ATP production and circadian cycling in these cells (**Fig. S5**). Together, these results provide evidence for a causal relationship between metabolic activity and circadian rhythmicity in PDA, and suggest that modulation of OxPhos is capable of strengthening circadian cycling in these cells only if it establishes a hypometabolic state.

### Circadian cycling strength varies with metabolic state among human patient-derived melanoma cell lines

Modulation of circadian cycling strength in PDA cell lines through pharmacological manipulation of OxPhos provides causal evidence that the circadian clock is inhibited by metabolic hyperactivity in a mouse cancer model. However, questions remained about whether this association would be observable across a wider range of cancer cell lines as well as its potential relevance for human disease. To investigate the association between metabolic activity and circadian rhythms in these contexts, we generated Bmal1::luciferase reporter lines from a library of human patient-derived melanoma lines and compared their circadian and metabolic activity (**Fig. 5, Fig. S6**, and **Table S1**) [26, 27]. For the analysis, we excluded lines that displayed arrhythmic profiles of the Bmal1::luciferase reporter, and one that had very high amplitude cycling, well outside the range seen in other lines (Z-score > 2). Among the lines that demonstrated robust circadian cycling, we observed a negative association between Bmal1::luciferase oscillation amplitude and both OxPhos-associated ATP production and basal glycolysis (**Fig. 5D** left and middle panels). In contrast, we did not observe any association between metabolic activity and circadian period in these cell lines (**Fig. S7**). Interestingly, melanoma lines that had high levels of ATP production also tended to have high levels of basal glycolysis, suggesting that these cell lines are distributed along an axis between hypometabolic and hypermetabolic activity. These results demonstrate that hypermetabolic activity is associated with circadian rhythm disruption in patient-derived melanoma cells.

**Figure 5.**
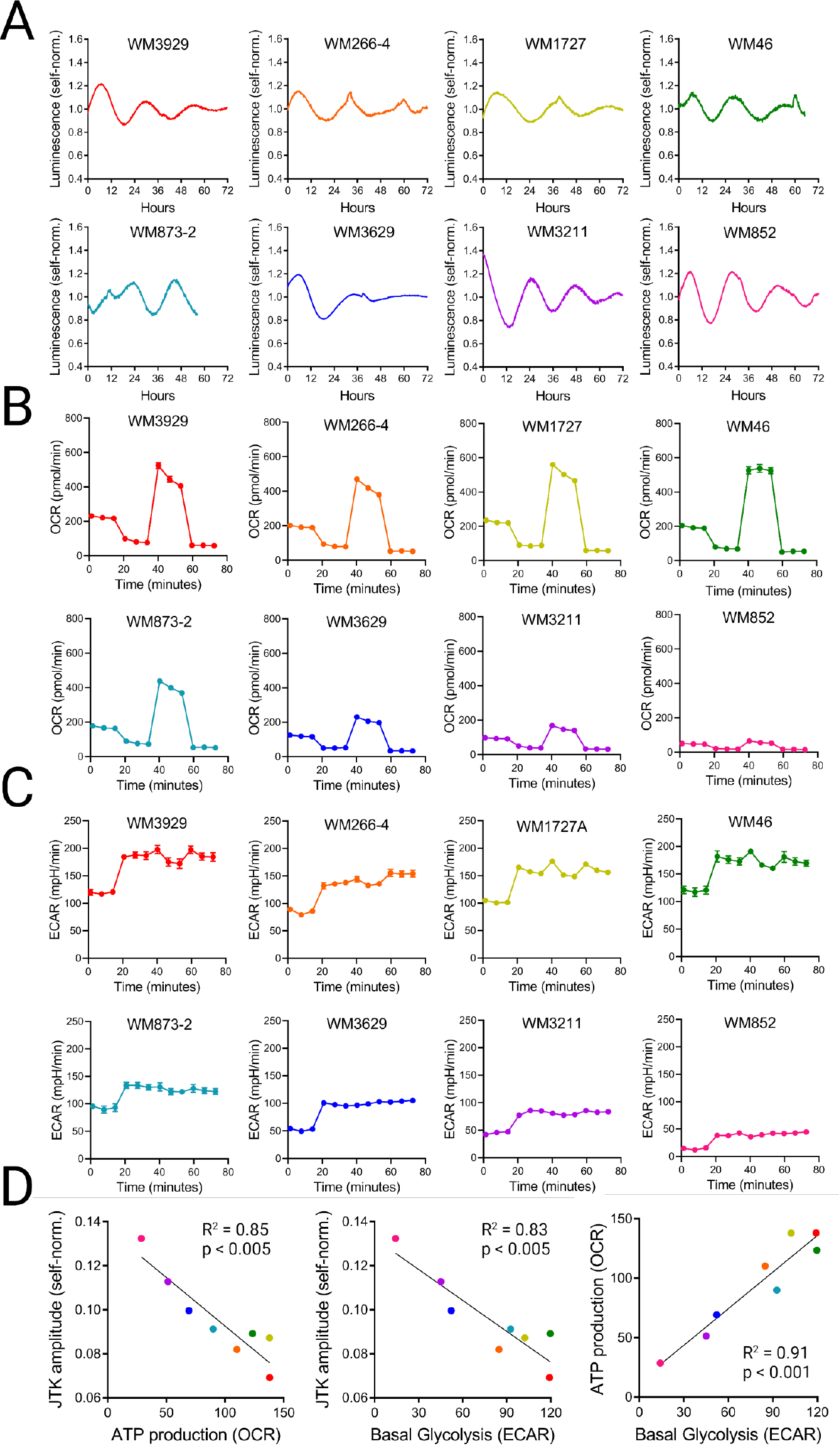
Circadian and metabolic phenotypes of human patient-derived melanoma lines. (**A**) Self-normalized luminescence traces of Bmal1::luciferase activity recorded from melanoma cell lines. (**B**) Oxygen consumption rates of melanoma cell lines during Seahorse mitochondrial stress test. (**C**) Extracellular acidification rates of melanoma cell lines during Seahorse mitochondrial stress test. (**D**) Linear correlations between JTK amplitude and ATP production (left), JTK amplitude and basal glycolysis (middle), and ATP production and basal glycolysis (right) across melanoma cell lines.

## Discussion

We report metabolic control of circadian cycling in PDA, such that robust circadian cycling is detected only in cancer cells with low levels of glycolysis and OxPhos. Additionally, among a panel of patient-derived melanoma lines, we observed that hypermetabolic lines tended to have weaker circadian oscillation. We infer that high metabolic activity is incompatible with maintenance of circadian cycling.

To explore the crosstalk between metabolism and circadian rhythms, we used the classical luciferase reporter system and realized that the luciferase dependence on ATP results in a simultaneous readout of OxPhos and circadian transcriptional activity. We show in circadian luciferase reporter MAFs that normalizing the luminescence of drug-treated groups to the control condition measures a combination of ATP production and circadian transcription, while normalizing each group to itself isolates the signal’s circadian component. As the mutual regulation between metabolism and circadian rhythms in health and disease remains a topic of significant interest, we strongly recommend carefully evaluating the use and interpretation of circadian luciferase reporter systems for each study.

Applying this system to a mouse model of PDA, we show considerable diversity among metabolic and circadian profiles from a panel of tumor cell clones. Across these cell lines we observed a strong inverse association between overall metabolic activity and the strength of circadian cycling. PDA lines with hypo- and hypermetabolic activity demonstrated the strongest and weakest cycling respectively, while cell lines whose metabolic profile suggested reliance on either glycolysis or OxPhos demonstrated intermediate circadian phenotypes. Inhibition of OxPhos was sufficient to enhance circadian cycling only in the single PDA line that presented with both high OxPhos and low glycolysis prior to treatment. Together these results suggest that glycolytic and OxPhos activity each contribute toward circadian dysfunction in PDA. Interestingly, expression of several clock genes is dysregulated in human PDA patient tumors and decreased expression of circadian transcription factor BMAL1 is a predictor of tumor progression, suggesting that metabolic reprogramming in PDA might drive cancer progression in part through dysregulation of the circadian clock [12, 28]. We further show that patient-derived melanoma cell lines are distributed along an axis between hypometabolic and hypermetabolic activity, and that this phenotype is strongly associated with the amplitude of circadian oscillations.

An intriguing set of candidates as effectors of glycolytic and OxPhos-mediated inhibition of circadian cycling is the sirtuin family of NAD-dependent deacetylases. Of the seven mammalian sirtuins, two (SIRT3 and SIRT1) localize to mitochondria and the cytoplasm respectively and function as integrators of metabolic activity that interact with the molecular clock [29-31]. In mitochondria, elevated consumption of NADH by complex I to produce excess NAD^+^ may decouple SIRT3 activity from the circadian clock, impairing its function as a mediator of circadian control over mitochondrial activity [6, 7]. This disruption of circadian mitochondrial activity might itself lead to dampened cycling amplitude by desynchronizing the metabolic pathways that reinforce transcriptional rhythms [9, 32]. Similarly, excess cytoplasmic NAD^+^ produced through conversion of pyruvate to lactate by lactate dehydrogenase in the context of elevated glycolysis may drive SIRT1 destabilization of BMAL1 and dampen circadian cycling [30, 33]. Notably, sirtuins themselves have also been linked to cancer progression, including in PDA, raising the possibility that these genes could present a potential therapeutic target [12, 34]. Another possibility is that buildup of lactate itself in the cytoplasm may inhibit circadian rhythmicity. Conversion of pyruvate to lactate drives intracellular acidification, which may disperse lysosomes away from the nucleus resulting in dysregulation of mTOR signaling and thereby disrupted circadian cycling [35].

Our results suggest a strong connection between metabolic state and circadian function in cancer. Knowing whether a particular cancer maintains circadian cycling is relevant for chronotherapeutic approaches, many of which target the timing of the cell cycle for optimal drug efficacy [36-39]. Future work to implicate the circadian clock as an effector of metabolically driven cancer progression may also create new avenues for cancer treatment. Overall, the link between metabolism and circadian rhythms we describe may be broadly relevant for disease states associated with inflammation-associated metabolic reprogramming including several forms of cancer, neurodegeneration, and stroke [40].

## Methods

### Cell Culture

PDA cells provided by B. Z. Stanger were cultured in Dulbecco’s Modified Eagle’s Medium (DMEM; 11995-065, Gibco) containing 10% fetal bovine serum (FBS; S11150, Atlanta Biologicals) and 1:100 antibiotic-antimycotic (15140-122, Gibco). For bioluminescent recording, cells were transferred to recording media containing 0.35% sodium bicarbonate (S5761, Sigma), 0.35% glucose (G7021, Sigma), 10mM HEPES (15630-080, Gibco), 10% FBS, and 0.2 mM luciferin (14681, Cayman Chemical) in DMEM (D-2902, Sigma). After the addition of recording media plates were sealed with TopSeal-A Plus plate covers (6050185, Perkin Elmer).

Patient-derived melanoma cells were cultured in MCDB 153 media (M6395, Millipore) containing 18% Leibovitz’s L-15 media (11415064, ThermoFischer), 2% fetal bovine serum, and 1.68 mM CaCl2. For bioluminescent recording, cells were transferred to recording media containing 0.5% sodium bicarbonate solution (S8761, Sigma), 10mM HEPES, 5% FBS, 1:400 antibiotic-antimycotic, 0.1 mM luciferin, and 100nM dexamethasone in RPMI 1640 (90022PB, Mediatech). After the addition of recording media plates were sealed with TopSeal-A Plus plate covers (6050185, Perkin Elmer). All cells were maintained at 37°C and 5% CO2 and were confirmed to be mycoplasma-free.

### Circadian rhythm reporter cell line generation

pPer2-dLuc-eGFP lentiviral reporter was provided by A. C. Liu at the Department of Physiology and Functional Genomics, University of Florida. Stable PDA cell reporter lines expressing Per2-dLuc and eGFP were generated according to stable transduction protocol using lentivirus-mediated gene delivery as previously described [41]. Briefly, LentiX 293T cells (632180, Clontech) were grown to 70% confluence in 10 cm dishes, and co-transfected with pPer2-dLuc-eGFP, together with the packaging plasmids (pDVPR8.1 and pVSV-G, Addgene) in a 10:1:0.5 ratio, using Lipofectamine 3000 (L3000008, Life Tech) according to the manufacturer’s directions. 24 hours following transduction, the >50% transfection efficiency was confirmed via fluorescent microscopy to assess GFP positivity, and the media was exchanged. Forty-eight hours following transduction, the supernatant was collected from the transfected 293T cells, and the supernatant was centrifuged at 1000 × g for 5 min to pellet any 293T cells. Finally, stable PDA cell lines were generated by infecting 70% confluent cultures with the recombinant lentiviral vectors with 10 mg/mL polybrene (TR-1003, Sigma-Aldrich) twice over 2 consecutive days, and then individual GFP^+^ cells were sorted on the FACSMelody (BD Biosciences) to establish clonal reporter lines.

### Mitochondrial Stress Test

The mitochondrial function of MAFs was measured with a Seahorse XF Cell Mito Stress Test Kit (103015-100, Agilent) using a Seahorse XF96 Extracellular Flux Analyzer. A total of 1.5 × 10^5^ Bmal1::luciferase MAFs or Per2::luciferase PDA cells were seeded per well in 96-well black-sided imaging plates (101085-004, Agilent) pre-coated with 0.2% gelatin for 1 hour at 37°C. Cells were treated with DMSO, BEZ (500 μM), ROT (20 nM), or DMF (60 μM) for 48 hours prior to the assay. To measure the oxygen consumption rate and extracellular acidification rate of MAFs, mitochondrial complex inhibitors (oligomycin 1 μM, FCCP 0.5 μM, rotenone/antimycin A 1 μM) were successively added to the cell culture microplate to measure key parameters of mitochondrial function with the Seahorse XF96 Analyzer. To measure the oxygen consumption rate and extracellular acidification rate of patient-derived melanoma cell lines, concentrations of 1.5 μM oligomycin, 1 μM FCCP, and 1 μM rotenone/antimycin A were used. Each condition was assayed in a minimum of five replicates per experiment.

### Metabolic Profiling

#### Metabolomic Extraction

Metabolomic extraction from cells was done as described [42]. 1 mL of cold 80% MeOH from −80°C, 40 μL of Metabolomics ISTD mix were added to each plate. Cells were scraped and transferred to microcentrifuge tubes in ice. Samples were pulse-sonicated in ice with a sonic dismembranator (Fisher Scientific) for 30 sec, incubated on ice for 10 min, and then pulsed again for 30 sec. Samples were pelleted by centrifugation at 6000 x g for 5 min at room temperature. 500 μL of supernatant was moved to a clean microcentrifuge tube, dried under nitrogen, and resuspended in 50 μL of 5% (w/v) SSA in water. 3 μL injections were used for LC-HRMS analysis.

#### Metabolomic LC-HRMS

Metabolites were separated using a XSelect HSS C18 column (2.1 mm x 150 mm, 3.5 μm particle size) (Waters, Milford, MA) in an UltiMate 3000 quaternary UHPLC (Thermo Scientific) equipped with a refrigerated autosampler (5°C) and column heater (50°C). Solvent A consisted of water with 5 mM DIPEA and 200 mM HFIP and Solvent B consisted of MeOH with 5 mM DIPEA and 200 mM HFIP. Flow gradient conditions were as follows: 0% B for 6 min at 0.18 mL min−1, increased to 1% B for 2 min at 0.2 mL min−1, increased to 2% B for 4 min, increased to 14% B for 2 min, increased to 70% B for 2 min, increased to 99% B for 1 min, increased flow rate to 0.3 mL min−1 for 0.5 min, increased flow rate to 0.4 mL min−1 for 4 min, then washed by decreasing to 0% B for 2.3 min at 0.3 mL min−1, decreased to 0.2 mL min−1 for 0.2 min, and ending with flow of 0.18 mL min-1. Samples were analyzed using a Q Exactive HF (QE-HF) (Thermo Scientific) equipped with a heated electro-spray ionization (HESI) source operated in the negative ion mode. Column effluent was diverted to the QE-HF from 0.5 to 19 min and then to waste for the remaining time of the run.

#### Bioluminescence recording and data analysis

A total of 1 × 10^6^ Bmal1::luciferase MAFs or Per2::luciferase PDA cells were seeded per well into 24-well black-sided imaging plates (1450-606, Perkin Elmer). Cells were treated with DMSO, BEZ (500 μM), ROT (20 nM), or DMF (60 μM) for 24 hours prior to recording. After the 1 μM dexmethasone pulse in DMEM (∼60 min) (D2915, Sigma-Aldrich), cells were transferred to bicarbonate recording media containing 0.2 mM luciferin and either DMSO, BEZ (500 μM), ROT (20 nM), or DMF (60 μM). Real-time bioluminescence of the cells was monitored using a LumiCycle luminometer (Actimetrics).

For patient-derived melanoma cells, 4 × 10^4^ cells expressing Bmal1::luciferase were seeded per well into 24-well black-sided imaging plates. Cells were incubated at 37°C and 5% CO2 for 2 days until they become confluent. On the day of the experiment, cells were transferred to recording media and the plate was sealed with adhesive optical PCR plate film before recording in a LumiCycle luminometer. DMSO-normalization of luminescence traces was performed by dividing each time point by the 24-hour rolling average of the DMSO condition luminescence for each experiment (T0 represents the first time point 12 hours after recording beings). Self-normalization of luminescence traces was performed by dividing each time point by the 24-hour rolling average of each independent trace. Circadian amplitude and period analysis of the luminescence data was performed using the JTK_CYCLE algorithm within the MetaCycle R package [43, 44].

#### RNA extraction and reverse transcription and quantitative PCR

A total of 1 × 10^6^ Bmal1::luciferase MAFs were seeded per 35-mm dish pre-coated with 0.2% gelatin for 1 hour at 37°C. Cells were treated with BEZ (500 μM), ROT (20 nM), or DMF (60 μM) for 12 hours prior to a 1 μM dexamethasone pulse (∼60 min). After 36 hours following dexamethasone synchronization, cells were detached with trypsin (25300-054, Gibco) and then pelleted. Total RNA was isolated from MAFs using the RNeasy Plus Mini Kit (74134, Qiagen) according to the manufacturer’s protocol. Equal amounts of complementary DNA were synthesized using the Invitrogen Superscript Vilo Master Mix (11755050, Life Technologies). qPCR was performed using the Taqman Gene expression PCR Master Mix (4369016, Applied Biosystems). All qPCR reactions were conducted at 50°C for 2 min, 95°C for 10 min, and then 40 cycles of 95°C for 15 s and 60°C for 1 min. The specificity of the reaction was assessed by melt curve analysis. The relative gene expression of each sample was quantified using the comparative Ct method. Samples were normalized to *Actin*, which was used as an endogenous control for all experiments. Experiments were performed using a ViiA7 Real-Time PCR machine (Thermo Fisher Scientific).

#### Cell Division and Apoptosis Assays

A total of 2 × 10^5^ Bmal1::luciferase MAFs or Per2::luciferase PDA cells were seeded per well in 96-well black-sided imaging plates (4517, Corning) pre-coated with 0.2% gelatin for 1 hour at 37°C. For the cell division assay, cells were treated with DMSO, BEZ (500 μM), ROT (20 nM), or DMF (60 μM) for 24 hours prior to the addition of 20 µM EdU and fixed 24 hours later. EdU staining was performed with the Click-iT EdU Cell Proliferation Kit (C10337, Invitrogen) according to the manufacturer’s instructions. For the apoptosis assay, cells were treated with DMSO, BEZ (500 μM), ROT (20 nM), or DMF (60 μM) for 48 hours prior to fixation. TUNEL staining was performed with the Click-iT Plus TUNEL Assay Kit (C10617, Invitrogen) according to the manufacturer’s instructions. Cells in all experiments were counterstained with 1:2000 Hoechst 3342 (6249, Thermo Scientific) to label nuclei.

#### Quantification and Statistical Analysis

All statistical tests used in this study were completed with Prism7 GraphPad software. For making multiple comparisons, we used one-way analysis of variance (ANOVA) followed by either Holm-Sidak’s multiple comparisons test to compare independent conditions or Benjamini, Krieger, and Yekutieli’s two-stage step-up procedure to control the false discovery rate when comparing multiple treatment conditions to a single control condition (p < 0.05).

Quantification of EdU and TUNEL Click-iT staining was performed with the Analyze Particles plugin within FIJI image analysis software. All DAPI and GFP images were background-subtracted and a threshold was applied to create a binary mask separating foreground from background. Cell nuclei and TUNEL particles were counted following application of a watershed algorithm to separate adjacent cells. ROIs were excluded from analysis if they contained fewer than 100 DAPI^+^ nuclei.

## Acknowledgements

We thank the Translational Biomarker Core at the University of Pennsylvania for their support with LC-HRMS experiments. This work was supported by grants from the NIH: F32MH125600 (to DMI), P30ES013508 (to CM), K99HL147212 (to SLZ), R01MH066912 (to SA), R01CA051497 (to CVD), R37NS048471 (to AS), and from the Howard Hughes Medical Institute (to AS). Figures were generated using BioRender.

## Contributions

DMI, CVD, and AS planned and designed the project. DMI performed and directed experiments, processed and analyzed data, and wrote the manuscript. XZ generated luciferase reporter patient melanoma lines and performed luciferase recording experiments. PB performed seahorse experiments with patient melanoma lines. CM, YS and SH performed LC-HRMS experiments and analyzed data. SLZ performed FAC sorting to isolate clonal luciferase reporter PDA cell lines. KC and PP performed qPCR experiments. AS managed the project and edited the manuscript with contributions from all co-authors.

## Notes

### Competing Interest Statement

The authors have declared no competing interest.

