## Supplemental Materials for "Hypermetabolic state is associated with circadian rhythm disruption in mouse and human cancer cells"

Supplementary Figure 1

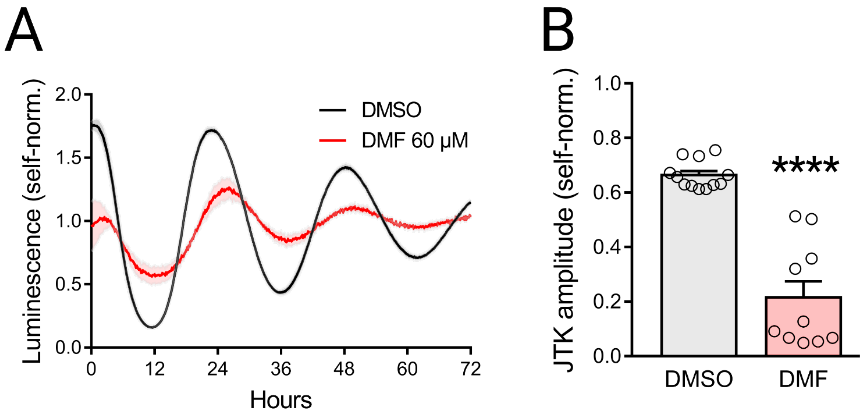


**Supplementary Figure 1. Nrf2 agonist DMF suppresses circadian rhythms in MAFs** (**A**) Self-normalized luminescence traces recorded from Bmal1::luciferase MAFs treated with DMF (n = 3 experiments). (**B**) Oscillation amplitudes of traces from (A). Error bars indicate mean ± SEM. ****$p < 0.0001$, Student's t-test.

Supplementary Figure 2

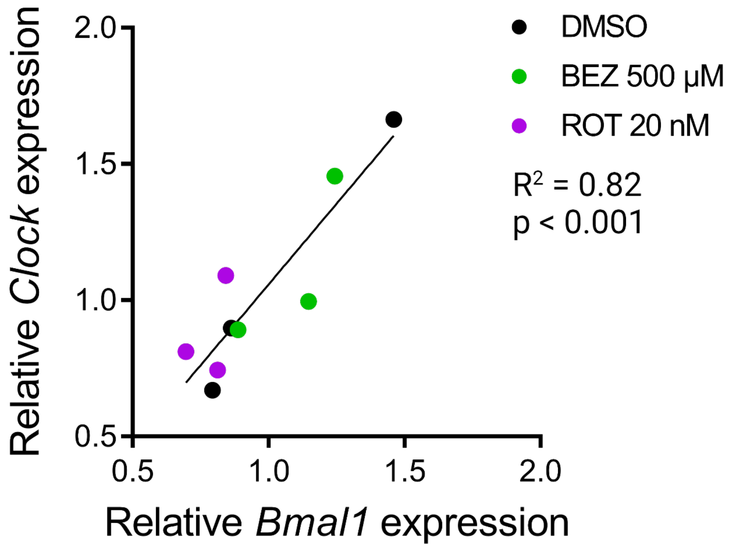


**Supplementary Figure 2. Association between *Bmal1* and *Clock* in MAFs** Linear correlation between peak *Bmal1* and *Clock* mRNA expression across 48-hour drug treatment groups measured with qPCR relative to *Actin*.

Supplementary Figure 3

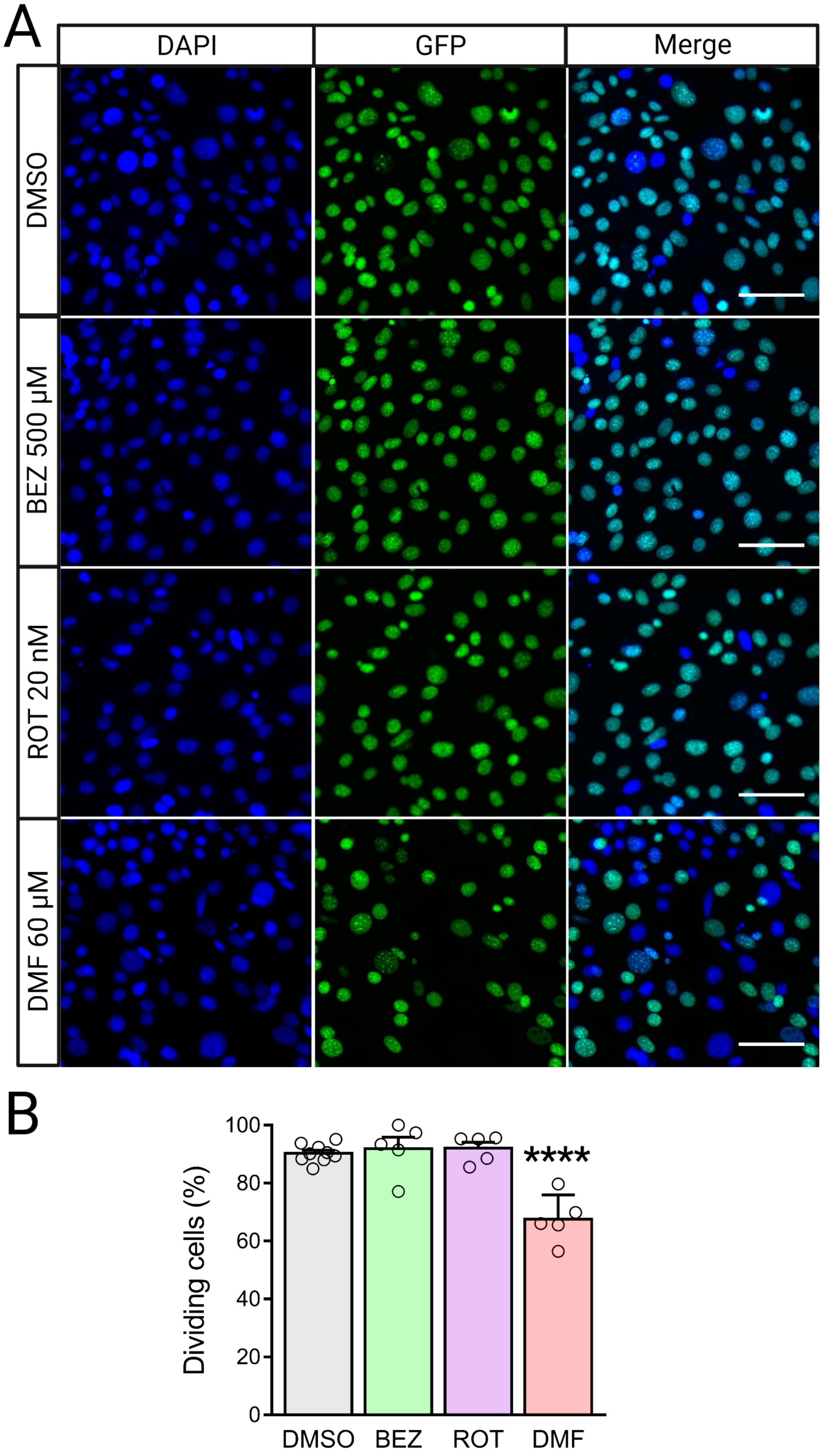


**Supplementary Figure 3. MAF cell division following drug treatment** (**A**) Representative images of DAPI and EdU nuclei staining in MAF cultures following 48 hours of treatment with BEZ, ROT, or DMF (**B**) Percentage of $EdU^+$ nuclei (dividing cells) from MAF cultures following drug treatment (n = 6-9 ROIs from 2-3 biological replicates). Scale bars: 100 µm. Error bars indicate mean ± SEM. ****$p < 0.0001$, one-way ANOVA test with Benjamini, Krieger, and Yekutieli's two-stage step-up procedure to control the FDR, drug treatment conditions compared to DMSO control.

Supplementary Figure 4

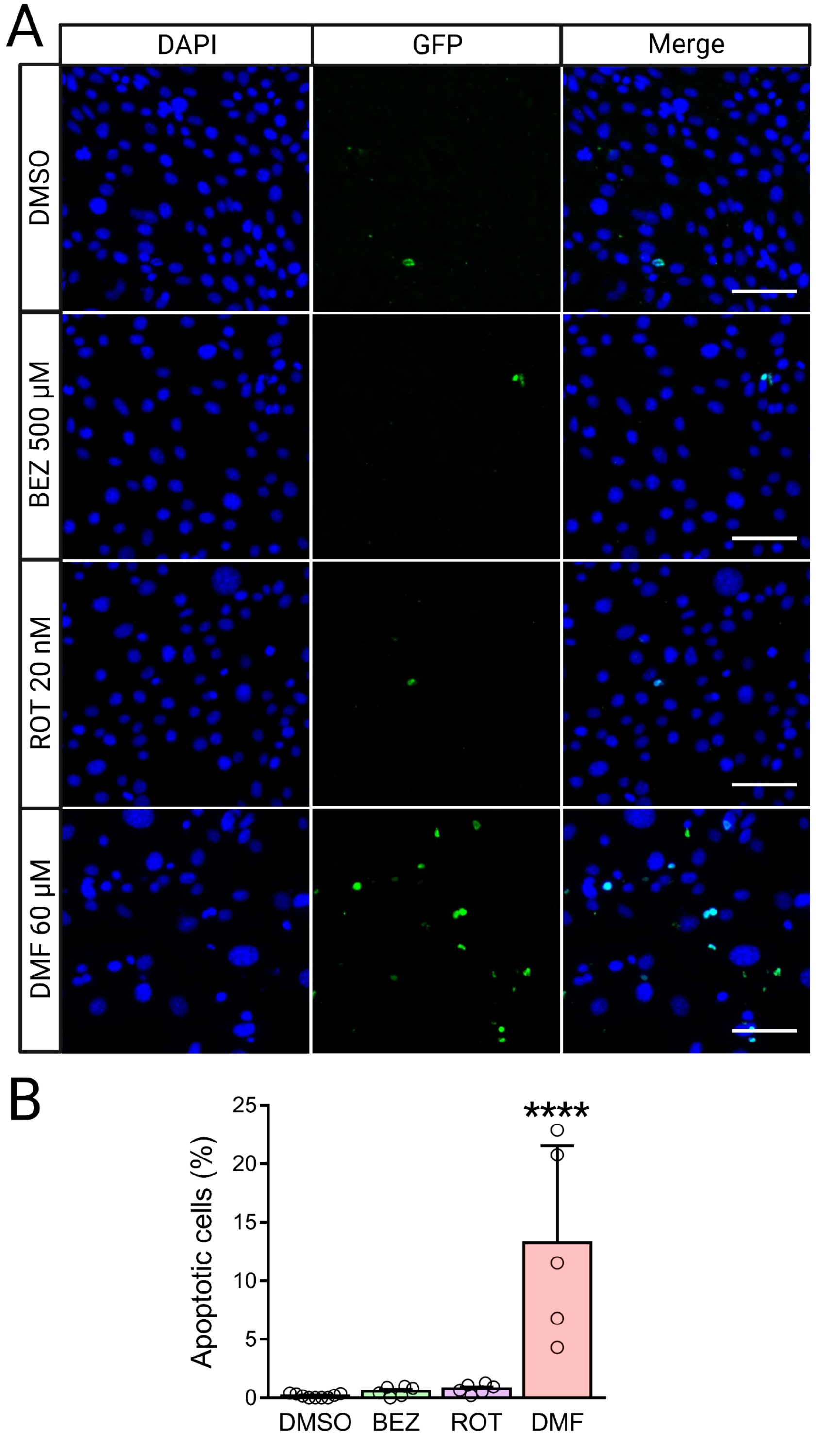


**Supplementary Figure 4. MAF apoptosis following drug treatment** (**A**) Representative images of DAPI and TUNEL staining in MAF cultures following 48 hours of treatment with BEZ, ROT, or DMF (**B**) Percentage of $TUNEL^+$ particles (apoptotic cells) from MAF cultures following drug treatment (n = 6-9 ROIs from 2-3 biological replicates). Scale bars: 100 µm. Error bars indicate mean ± SEM. ****$p < 0.0001$, one-way ANOVA test with Benjamini, Krieger, and Yekutieli's two-stage step-up procedure to control the FDR, drug treatment conditions compared to DMSO control.

Supplementary Figure 5

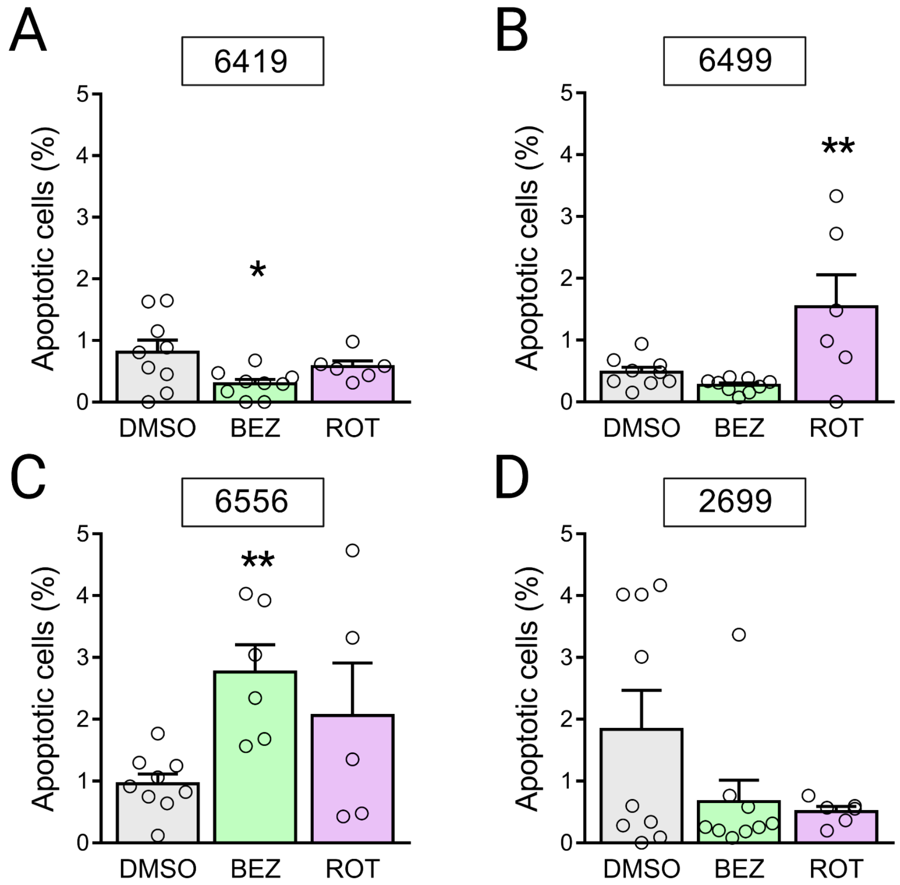


**Supplementary Figure 5. PDA apoptosis following metabolic drug treatment** (**A-D**) Percentage of TUNEL$^+$ cells from PDA cell lines 6419 (A), 6499 (B), 6556 (C), and 2699 (D) (n = 6-9 ROIs from 2-3 biological replicates). Error bars indicate mean ± SEM. *$p < 0.05$, **$p < 0.01$, one-way ANOVA test with Benjamini, Krieger, and Yekutieli's two-stage step-up procedure to control the FDR, drug treatment conditions compared to DMSO control.

Supplementary Figure 6

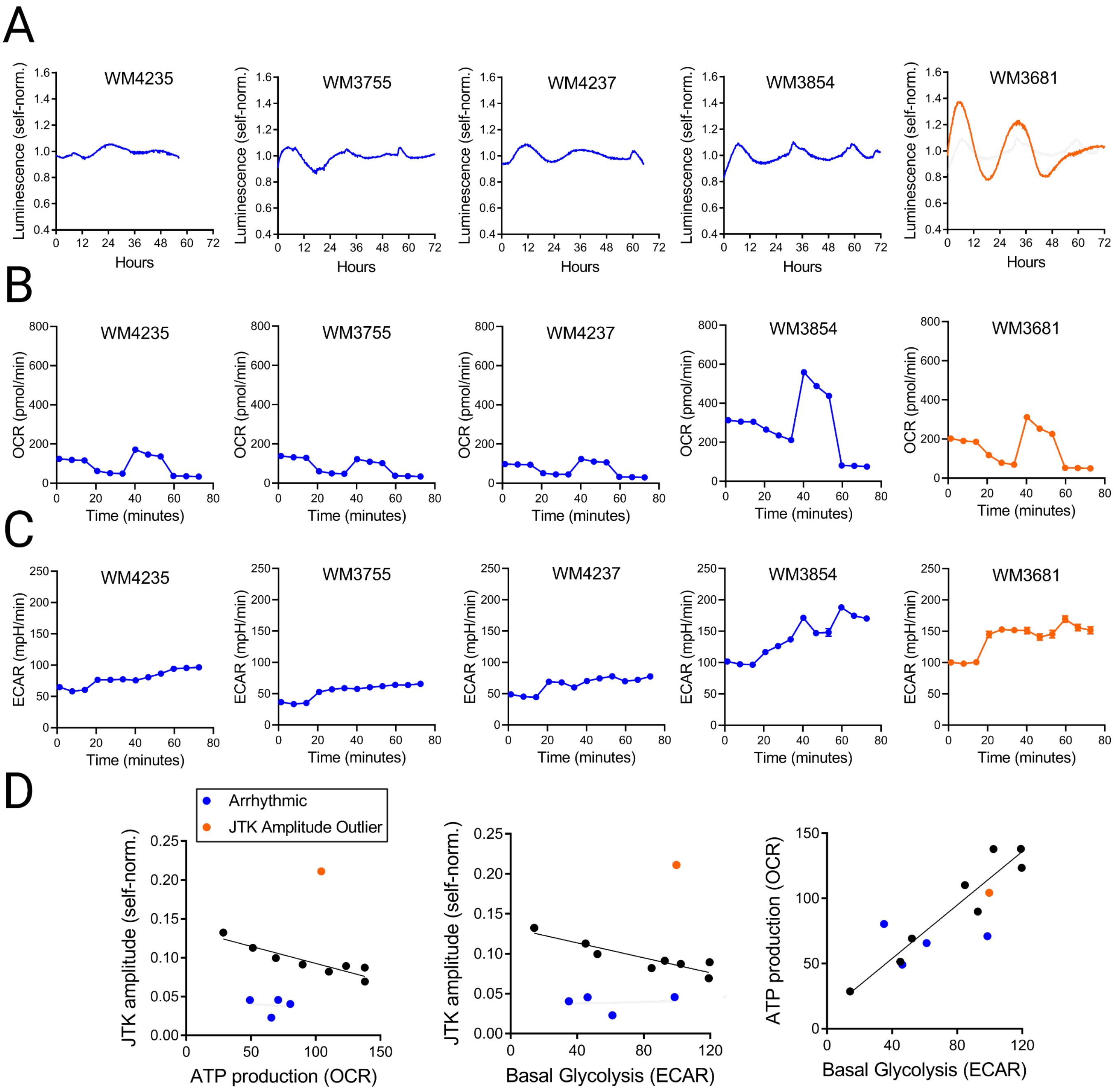


**Supplementary Figure 6. Circadian and metabolic phenotypes of excluded patient-derived melanoma lines** (**A**) Self-normalized luminescence traces ± SEM of Bmal1::luciferase activity recorded from melanoma cell lines excluded due to arrhythmicity (blue) or outlier cycling amplitude (orange). (**B**) Oxygen consumption rates of melanoma cell lines during Seahorse mitochondrial stress test. (**C**) Extracellular acidification rates of melanoma cell lines during Seahorse mitochondrial stress test. (**D**) Excluded melanoma lines plotted relative to linear correlations between JTK amplitude and ATP production (left), JTK amplitude and basal glycolysis (middle), and ATP production and basal glycolysis (right) across melanoma cell lines. Error bars indicate mean ± SEM.

Supplementary Figure 7

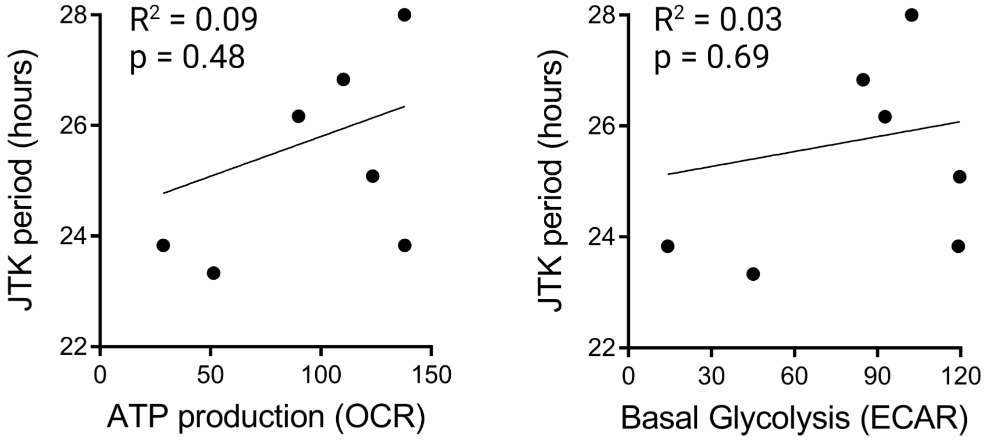


**Supplementary Figure 7. No association between circadian period and metabolic phenotypes across patient-derived melanoma cell lines** Linear correlations between JTK period (hours) and either ATP production (left) or basal glycolysis (right) across melanoma cell lines.

### Supplementary Table 1

| Patient-Derived Melanoma Line | JTK Amplitude | JTK Period (hours) | ATP Production (OCR) | Basal Glycolysis (ECAR) | Exclusion Criteria |
|---|---|---|---|---|---|
| WM3929 | 0.069 | 23.83 | 137.99 | 119.16 | N/A |
| WM266-4 | 0.082 | 26.83 | 110.25 | 84.78 | N/A |
| WM1727 | 0.087 | 28.00 | 137.88 | 102.32 | N/A |
| WM46 | 0.089 | 25.08 | 123.46 | 119.63 | N/A |
| WM873-2 | 0.091 | 26.17 | 89.90 | 92.71 | N/A |
| WM3629 | 0.100 | 28.58 | 69.20 | 52.18 | N/A |
| WM3211 | 0.113 | 23.33 | 51.49 | 45.03 | N/A |
| WM852 | 0.132 | 23.83 | 28.68 | 14.16 | N/A |
| WM4235 | 0.023 | 35.17 | 65.75 | 61.20 | Arrhythmic |
| WM3755 | 0.041 | 24.17 | 80.43 | 35.01 | Arrhythmic |
| WM4237 | 0.045 | 27.50 | 49.15 | 46.26 | Arrhythmic |
| WM3854 | 0.046 | 26.25 | 71.03 | 98.60 | Arrhythmic |
| WM3681 | 0.211 | 28.50 | 104.28 | 99.69 | JTK Amplitude Outlier |

**Supplementary Table 1. Circadian and metabolic characteristics of patient-derived melanoma cell lines** Oscillation amplitudes and periods were calculated with JTK analysis of self-normalized Bmal1::luciferase recordings. ATP production (OCR; pmol/min) was calculated as the difference in oxygen consumption rate before and after treatment with the ATP synthase inhibitor oligomycin. Basal Glycolysis (ECAR; mpH/min) was calculated as the baseline extracellular acidification rate of melanoma lines prior to oligomycin treatment. Arrhythmic cell lines were defined as having JTK amplitudes less than 0.005. Cell lines with JTK amplitude Z-scores greater than 2 were excluded from analysis.
